# Decoding grasp and speech signals from the cortical grasp circuit in a tetraplegic human

**DOI:** 10.1101/2021.10.29.466528

**Authors:** S. K. Wandelt, S. Kellis, D. A. Bjånes, K. Pejsa, B. Lee, C. Liu, R. A. Andersen

## Abstract

Tetraplegia from spinal cord injury leaves many patients paralyzed below the neck, leaving them unable to perform most activities of daily living. Brain-machine interfaces (BMIs) could give tetraplegic patients more independence by directly utilizing brain signals to control external devices such as robotic arms or hands. The cortical grasp network has been of particular interest because of its potential to facilitate the restoration of dexterous object manipulation. However, a network that involves such high-level cortical areas may also provide additional information, such as the encoding of speech. Towards understanding the role of different brain areas in the human cortical grasp network, neural activity related to motor intentions for grasping and performing speech was recorded in a tetraplegic patient in the supramarginal gyrus (SMG), the ventral premotor cortex (PMv), and the somatosensory cortex (S1). We found that in high-level brain areas SMG and PMv, grasps were well represented by firing rates of neuronal populations already at visual cue presentation. During motor imagery, grasps could be significantly decoded from all brain areas. At identical neuronal population sizes, SMG and PMv achieved similar highly-significant decoding abilities, demonstrating their potential for grasp BMIs. During speech, SMG encoded both spoken grasps and colors, in contrast to PMv and S1, which were not able to significantly decode speech.These findings suggest that grasp signals can robustly be decoded at a single unit level from the cortical grasping circuit in human. Data from PMv suggests a specialized role in grasping, while SMG’s role is broader and extends to speech. Together, these results indicate that brain signals from high-level areas of the human cortex can be exploited for a variety of different BMI applications.

## Introduction

The ability to grasp and manipulate everyday objects is a fundamental skill, required for most daily tasks of independent living. Functional loss of this ability, due to partial or complete paralysis from a spinal cord injury (SCI), can irrevocably degrade an individual’s autonomy. People with tetraplegia have consistently rated recovery of hand and arm function as the highest priority for increasing their quality of life (Anderson, 2004),(Snoek et al., 2004). Brain-machine interfaces (BMI) could give tetraplegic individuals greater independence by directly recording neural activity from the brain and decoding these signals to control external devices such as a robotic arm or hand (Aflalo et al., 2015). Intracortical BMI’s use microelectrode arrays to capture the action potentials of individual neurons with a high signal to noise ratio (SNR) and high spatial resolution (Nicolas-Alonso and Gomez-Gil, 2012). If placed in brain areas of the grasp circuit in human, these devices are well suited to extract the neuronal signals supporting the control of a high-dimensional prosthetic hand (Collinger et al., 2013).

In this work, we will evaluate the encoding of grasp motor imagery in human supramarginal gyrus (SMG), a sub region of the posterior parietal cortex (PPC), the ventral premotor cortex (PMv) and the primary sensory cortex (S1). These brain areas are key components of the cortical grasp circuit. PPC and PMv each process complex cognitive processes, like goal and end-target directed signals (Aflalo et al., 2015), and low level trajectory and joint-angle motor commands, as M1 (Andersen et al., 2014), (Schaffelhofer and Scherberger, 2016). Decoding movement intentions from upstream brain areas such as PPC and PMv, instead of decoding individual finger movements from M1, may allow for more rapid and intuitive control of a grasp BMI (Andersen et al., 2014),(Andersen et al., 2019). S1 processes incoming sensory feedback signals from the peripheral nervous system. While it is not thought to participate in grasp planning per se, it processes proprioceptive signals during movement (Goodman et al., 2019), which could be exploited for a grasp BMI.

The grasp circuit was first identified in a non-human primate model (NHP), a network of cerebral pathways involved from visual object presentation to grasp execution. During an object manipulation task, neurons in anterior intraparietal cortex (AIP), a sub-region of PPC, and F5 (a PMv analog in NHP) have both shown selectivity for object features, such as size, shape and orientation (Murata et al., 1997), (A. Murata et al., 2000),(Sakata, 1995),(Taira, 1998). In contrast to M1, information about the attempted grasp can already be decoded during cue presentation with high accuracy, implicating a role in motor planning (Carpaneto et al., 2011), (Townsend et al., 2011), (Michaels and Scherberger, 2017), (Schaffelhofer and Scherberger, 2016). In human electrophysiological studies, some of these results have been replicated in AIP, demonstrating the ability to decode grasp planning and intention while a human participant performed motor imagery of one of five cued grasps (Klaes et al., 2015). However, it remains to be seen if an analogous relationship exists between pre-clinical results in NHP F5 and human PMv.

The supramarginal gyrus (SMG) has been hypothesized as a region specialized for complex tool use, evolutionarily evolving from a duplicate region of NHP AIP (Orban and Caruana, 2014). Functional magnetic resonance imagining (fMRI) studies have shown activation of SMG during observed tool use, a finding which could not be replicated in presumed analogous anatomical regions of cortex in NHP (Peeters et al., 2009). Other studies confirmed SMG activity modulates during grasping and manipulation of objects (Sakata, 1995), reaching (Filimon et al., 2009), and tool use (Gallivan et al., 2013) (Orban and Caruana, 2014), (McDowell et al., 2018), (Reynaud et al., 2019). Additionally, one study demonstrated SMG’s involvement in both the planning and execution of (pantomimed) tool use (Johnson-Frey, 2004). These characteristics highlight SMG’s rich potential as a source of grasp related neural signals in human cortex, which could be exploited for BMI control of a prosthetic hand. Furthermore, transcranial magnetic stimulation (TMS) and fMRI studies have extensively documented SMG’s involvement in language processing (Stoeckel et al., 2009), (Sliwinska et al., 2012), (Oberhuber et al., 2016) and verbal working memory (Deschamps et al., 2014). This evidence suggests SMG could be involved in many high-level processes, such as speech, indicating its potential value for a variety of different BMI applications.

Recent studies in S1 indicate its potential as a target site for BMI applications in patients with tetraplegia. Human and NHP studies have demonstrated decoding of hand kinematics during executed hand gestures (Branco et al., 2017) and before contact during object grasping (Okorokova et al., 2020), respectively. These results suggest that neural signals could also be present during imagined movement (Zhang et al., 2017). Furthermore, modulation of S1 neurons during motor imagery of reaching has been demonstrated for the same participant whose data underlies this work (Jafari et al., 2020). If grasping motor imagery can be robustly decoded, a single implant in S1 could allow for a bidirectional BMI, able to decode grasp intentions and utilize electrical stimulation to evoke somatosensations (Armenta Salas et al., 2018).

In this work, a tetraplegic participant performed motor imagery of several different grasps, while neurophysiological responses were captured from three implant sites using recording microelectrode arrays, the supramarginal gyrus, the ventral premotor cortex and the primary sensory cortex. We evaluated the decodability of these imagined grasps in the context of evaluating suitability for BMI applications. To explore each regions’ role in other high-level cognitive tasks, the participant performed verbal speech of names of grasps, and names of colors. We hypothesized that grasp motor imagery would modulate activity in all three brain areas, while SMG would show higher neuronal modulation during speech than PMv and S1, due to its involvement in visual word recognition and phonological processing (Oberhuber, Cerebral Cortex, 2016).

## Methods

### Participant and Implants

A tetraplegic participant was recruited for an IRB- and FDA-approved clinical trial of a brain-machine interface and he gave informed consent to participate. The participant suffered a spinal cord injury at cervical level C5 two years prior to participating in the study. The targeted areas for implant were left ventral premotor cortex (PMv), supramarginal gyrus (SMG), and primary somatosensory cortex (S1). For more information about implant location, see (Armenta Salas et al., 2018). To identify exact implant sites within these regions, the participant performed imagined reaching and grasping tasks during fMRI, described in (Aflalo, Kellis et al., 2015). In November 2016, the participant underwent surgery to implant one 96-channel multi-electrode array (Neuroport Array, Blackrock Microsystems, Salt Lake City, UT) in SMG and PMv each, and two 7 × 7 sputtered iridium oxide film - tipped microelectrode arrays with 48 channels each in S1.

### Data collection

Recording began two weeks after surgery and continued one to three times per week. Data for this work were collected between 2017 and 2019. Broadband electrical activity was recorded from the NeuroPort arrays using Neural Signal Processors (Blackrock Microsystems, Salt Lake City, UT). Analog signals were amplified, bandpass filtered (0.3-7500 Hz), and digitized at 30,000 samples/sec. To identify putative action potentials, these broadband data were bandpass filtered (250-5000 Hz), and thresholded at - 4.5 the estimated root-mean-square voltage of the noise. Waveforms captured at these threshold crossings were then spike sorted by assigning each observation to a putative single neuron, and the rate of occurrence of each “unit”, in spikes/sec, are the data underlying this work. Units with firing rate <1.5 Hz were excluded from all analyses. To allow for meaningful analysis of individual datasets, recording sessions where high levels of noise prevented us from isolating more than three units on an array were excluded. This resulted in the removal of three PMv datasets. The rounded average number of recorded units per session was 55 +/- 17 for SMG, 12 +/- 9 for PMv, and 119 +/- 48 for S1.

### Experimental Task

We implemented a task that cued five different grasps with visual images taken from the “Human Grasping Database” (Feix et al., 2016) to examine the neural activity related to imagined grasps in SMG, PMv and S1. The grasps were selected to cover a range of different hand configurations and were labeled “Lateral”, “WritingTripod”, “MediumWrap”, “PalmarPinch”, and “Sphere3Finger” (**Figure 1**A).

**Figure 1.**
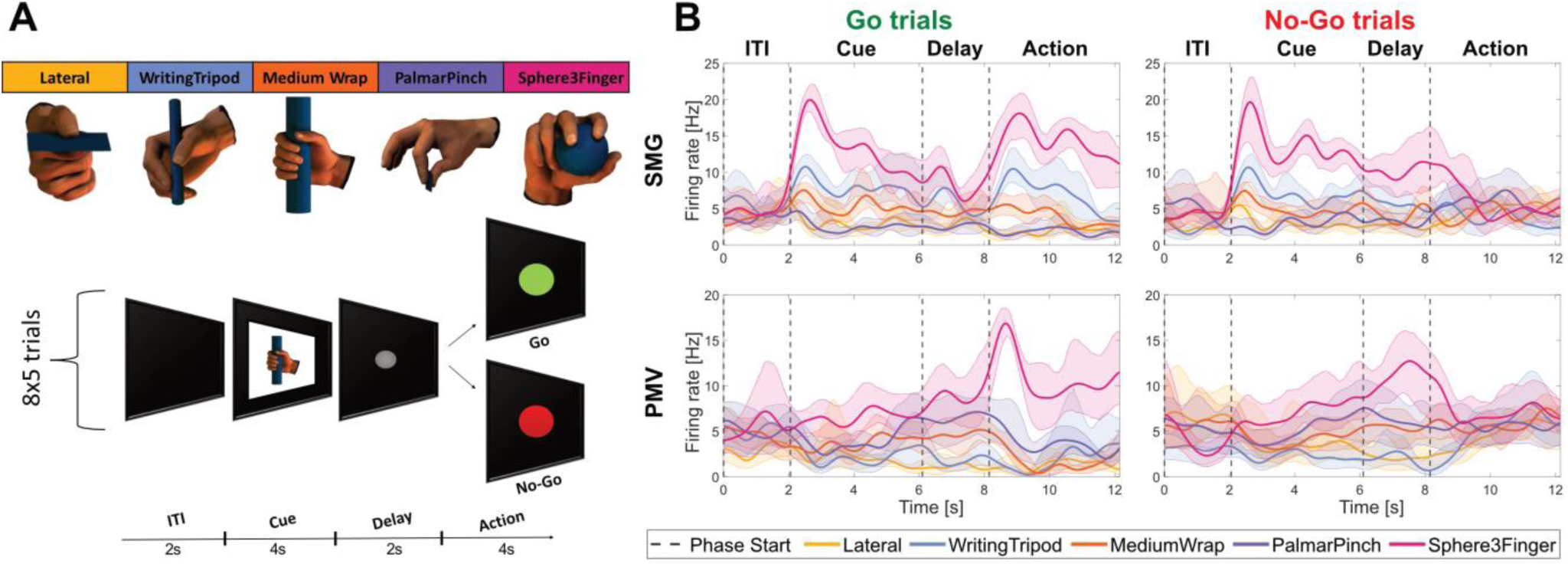
Neurons in posterior parietal cortex and ventral premotor cortex encode grasps. **A)** Grasp images from the “Human Grasping Database” *(Feix et al*., *2016)* were used to cue motor imagery in a tetraplegic human. The task was composed of an inter-trial interval (ITI), a cue phase displaying one of the grasp images, a delay phase and an action phase. The action phase was composed of intermixed Go trials (green), during which the participant performed motor imagery and No-Go trials (red), during which the participant rested. **B)** Example smoothed firing rates of neurons in SMG and PMv during Go (left) and No-Go (right) trials. The plots show the smoothed average firing rate of two example units (solid line, shaded area 95% bootstrapped c.i.) for 8 trials of each grasp, with vertical lines representing the beginning of each phase.

### Go task

Each trial consisted of four phases, referred to in this paper as ITI, cue, delay and action (**Figure 1**B). The trial began with a brief inter-trial interval (2 sec), followed by a visual cue of one of the five specific grasps (4 sec). Then, after a delay period (gray circle onscreen; 2 sec), the participant was instructed to imagine performing the cued grasp with his right (contralateral) hand (Go condition; green circle on screen; 4 sec). Three datasets had a longer action phase. For these, only data from the first four seconds of the action phase were included in the analysis.

### Go/No-Go task

In a Go/No-Go variation of this task, the participant was presented with either a green circle (Go condition) or a red circle (No-Go condition) after the delay, with instruction to imagine performing the cued grasp as normal during the Go condition, and to do nothing for the No-Go condition. In both variations of the task, conditions and grasp types were pseudorandomly interleaved and balanced with eight trials collected per combination (**Figure 1**B).

### Spoken Grasps task

A speaking variation of the task was constructed with the same task design outline above, but instead of performing motor imagery during the action phase, the participant was instructed to speak aloud the name of the grasp. **Spoken Colors task:** Another variation of this speaking task used five different colors instead of five grasps, and the participant was instructed to say the name of the color during the action phase (**Figure 5**A,B). On each session day, “Go task”, a “Spoken Grasps task” and a “Spoken Colors task” was performed, to allow comparisons between tasks.

Table 1 illustrated the number of recording sessions for each task variation.

**Table 1.** Number of recorded sessions for each task variation

| Task Area | Go task | Go/No-Go task | Spoken Grasps | Spoken Colors |
| --- | --- | --- | --- | --- |
| SMG | 6 | 9 | 5 | 5 |
| PMV | 6 | 6 | 5 | 5 |
| S1 | 6 | 7 | 5 | 5 |

The participant was situated 1 m in front of a LED screen (1190 mm screen diagonal), where the task was visualized. The task was implemented using the Psychophysics Toolbox (Brainard, 1997; Pelli, 1997; Kleiner et al, 2007) extension for MATLAB (MATLAB. (2018). 9.7.0.1190202 (R2019b). Natick, Massachusetts: The MathWorks Inc.).

### Neural Firing Rates

Firing rates of sorted units were computed as the number of spikes that occurred in 50ms bins, divided by the bin width, and smoothed using a Gaussian filter with kernel width of 50ms to form an estimate of the instantaneous firing rates (spikes/sec). For the Go condition, 40 trials (8 repetitions of 5 grasps) were recorded per block. For the No-Go condition, two consecutive blocks of 40 trials (4 repetitions of 5 Go and 5 No-Go grasps) were recorded and combined, to accommodate the participant with shorter tasks.

### Linear regression analysis

To identify units that exhibited selective firing rate patterns (or tuning) for the different grasps, linear regression analysis was performed in two different ways: 1) step by step in 50ms time bins to allow assessing changes in neuronal tuning over the entire trial duration 2) averaging the firing rate of specified time windows during cue (1.5s) and action phase (2s), allowing to compare tuning between both phases. The model returns a fit that estimates the firing rate of a unit based on the following variables:

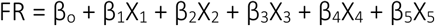

Where FR corresponds to the firing rate of that unit, and β corresponds to the estimated regression coefficients. A 48 × 5 indicator variable, X, indicated which data corresponded to which grasp. The first 8 rows were the average firing rate of the ITI phase, and indicated the offset term β_o,_ or baseline condition. These rows had only zeros. The next 40 rows indicated the trial data, for example, if the first trial was “Lateral” (grasp 1), it would have a 1 in column 1, and zeros in all other columns.

In this model, β symbolizes the change of firing rate from baseline for each grasp. A student’s t –test was performed to test the hypothesis of β = 0. A unit was defined as tuned if the hypothesis could be rejected (p < 0.05, t-statistic). This definition allows for tuning of a channel to zero, one, or multiple grasps during different time points of the trial.

### Linear regression significance testing

To assess significance of unit tuning, a null dataset was created by repeating linear regression analysis 1000 times with shuffled labels. Then, different percentile levels of this obtained null distribution were computed and compared to the actual data. Data higher than the 95th percentile of the null - distribution was denoted *, higher than 99th percentile was denoted **, and higher than 99.9th percentile was denoted ***.

### Classification

Using the neuronal firing rates recorded in this task, a classifier was used to evaluate how well the set of grasps could be differentiated during each phase. For each session and each array individually, naïve Bayes classification was performed, assuming an identical diagonal covariance matrix for each group. These assumptions, compared to a full diagonal covariance matrix, resulted in best classification accuracies. Classifiers were trained using averaged data from each phase, which were either 2s (ITI, delay) or 4s (cue, action). We applied principal component analysis (PCA) and selected the 10 highest principal components (PC’s), or PCs explaining more than 90% of the variance (whichever was higher), for feature selection on the training set. This feature selection method allowed us to compare if there was a correlation between the number of tuned units and classification accuracy, without selecting tuned units as features. The unit yield in PMv was generally lower than in SMG and S1, however, significant classification accuracies were still obtained with a limited number of features. Between 12 and 21 PCs were used in SMG, 6 and 16 in PMv, and 18 and 27 in S1. Leave one out cross-validation was performed to estimate decoding performance. A 95% confidence interval was computed by the student’s t inverse cumulative distribution function.

### Classification performance significance testing

To assess the significance of classification performance, a null dataset was created by repeating classification 1000 times with shuffled labels. Then, different percentile levels of this null distribution were computed and compared to the mean of the actual data. Mean classification performances higher than the 95th percentile were denoted *, higher than 99th percentile were denoted **, and higher than 99.9th percentile were denoted ***.

### Neuron dropping curve and cross phase classification

The neuron dropping curve represents the evolution of the classification accuracy based on the number of units used to train and test the model. All available units were used for all brain areas. Simultaneously to the neuron dropping analysis, cross-phase classification was performed to investigate how well a model trained on data of the cue phase can predict data of the action phase, and vice-versa. Classification with eightfold cross validation was performed for each subset of units (or features) selected for classification. First, one of the features was randomly selected, and the classification accuracy on cue and action phase was computed with a model trained on either the action phase or the cue phase. Then, a new subset of two random features was selected, and classification accuracy was again computed. This was performed until all features were added. PCA was performed on the dataset. To avoid overfitting by using more features than observations (40), the maximum number of principal components used was 20. The process was repeated 100 times, and the average prediction accuracy with 95% confidence interval (bootstrapping) was plotted against the number of neurons.

## Results

Grasp representation in SMG, PMv and S1 was characterized by implementing a task that cued a human participant to imagine five different grasps with visual images taken from the “Human Grasping Database” (Feix et al., 2016) (**Figure 1**A). The task contained four phases: an inter-trial interval (ITI), a cue phase, a delay phase, and an action phase, during which the participant performed motor imagery. In the Go variation of the task, motor imagery was performed during each action phase. In a Go/No-Go variation of the task, the action phase was composed of randomly intermixed Go trials and No-Go trials resulting in 10 experimental conditions (5 grasps × 2 action phases). This control allowed to verify that activity during the action phase was motor-related, and the participant can control motor imagery at will. We evaluated the brain regions’ potential for grasp BMI in two ways; firstly, by quantifying grasp tuning in the neuronal population and secondly, by assessing how well individual grasps were decodable from each area. We found that if large enough neuronal populations are present, both SMG and PMv show high grasp selectivity, making them noteworthy candidates for grasp BMI implantation sites. We also evaluated each region’s role during language processing. While our data indicates PMv is selectively active during grasping, SMG is highly engaged during speech production, pointing towards its potential application in a speech BMI.

As results were quantitatively similar during Go trials in both the Go and Go/No-Go variation of the task, neurons were pooled over both tasks for all session days (see Table 1), resulting in 819 SMG Go units, 504 SMG No-Go units, 146 PMv Go units, 78 PMv No-Go units, 1551 S1 Go units, and 948 S1 No-Go units.

Smoothed firing rates of example units for SMG and PMv during the Go/No-Go version of the task displayed neuronal modulation to the grasp “Sphere3Finger” in **Figure 1**B. Motor imagery evoked a strong response during the action phase of Go trials compared to the action phase of No-Go trials, where firing rate decreased back to baseline activity. This example SMG unit also showed an increase of firing rate during cue presentation, while the example PMv unit only showed an increase during action phase, providing examples of neurons previously labeled from NHP recordings studies as visuo-motor (tuned during both cue and action phase) and motor unit (tuned only during action phase) (Akira Murata et al., 2000), (Carpaneto et al., 2011),(Klaes et al., 2015).

After establishing individual neural firing rate modulation during motor imagery of different grasps, we quantified the entire neuronal population’s selectivity for each grasp. To compare selective neural activity within task stimuli presentation (image cue, Go-trial action phase, No-Go trial action phase), we determined the duration of selective (or tuned) activity of the neural population during each phase. In this context, tuning of a neuron to a grasp was determined by fitting a linear regression model to the firing rate in 50ms time bins. The p value associated with the coefficient estimate to each grasp was computed, and units that had significant p-values (< 0.05) to at least one grasp were defined as tuned.

### SMG, PMv and S1 show significant tuning to grasp during motor imagery

Population analysis revealed two main peaks of activation, one at the cue presentation and another during action phase (**Figure 2**A). During Go trials, the highest percentage of tuned units during cue presentation was in SMG (54.8%), achieving its peak 50ms earlier than PMv (41.1%). A decreased but sustained activity during the delay phase then led to a new rise during the action phase (39.0% for PMv, 37.4% for SMG). For S1, a minor increase in the number of tuned units was observed during action phase, but not during cue presentation.

**Figure 2.**
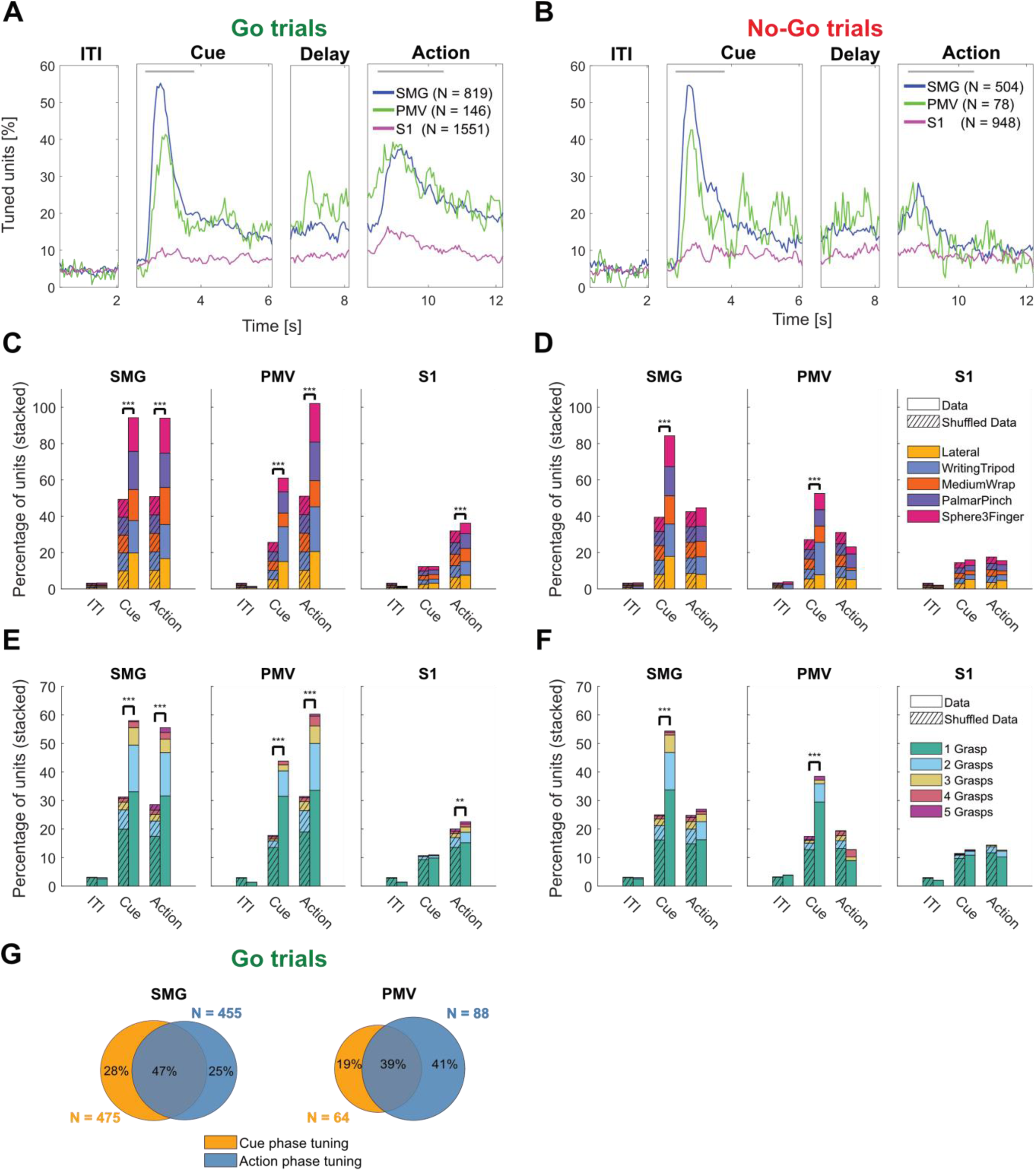
Population analysis of grasp tuning. **A)** Percentage of tuned channels to grasps for Go trials in 50ms time bins in SMG, PMv and S1 over the trial duration. The gray lines represent cue and action analysis windows for figures B,C and D. **B)** Same as A) for No-Go trials. **C)** Stacked percentage of units tuned for each grasp in ITI, cue phase and action phase window during Go trials. Significance was calculated by comparing data (right bar) to a shuffle distribution (striped lines, left bar). **D)** Same as C) for No-Go trials. **E)** Stacked percentage of units tuned to one, two, three, four and five grasps during cue phase and action phase analysis window during Go trials. Significance was calculated as described previously. **F)** Same as E) for No-Go trials. **G)** Overlap of tuned units between cue and action analysis window during Go trials for SMG and PMv.

During No-Go trials, neuronal activity peaked similarly during cue and delay phase, but then decreased around 1s after start of the action phase. This increase could indicate the formation of a motor plan during the cue phase, and a brief period of activity during the action phase when this plan was canceled (**Figure 2**B, No-Go trials, action phase).

The identified peaks of activity were selected to compute individual grasp tuning. Time windows incorporating the peaks began 250ms after the start of either cue or action phase (to account for processing latencies), and were respectively 1.5s and 2s long (gray lines, top of **Figure 2**A,B). Unlike the onset of the visual cue during the cue phase, it is not possible to measure the exact start time of motor imagery during action phase. Thus a longer time window (2s vs 1.5s) was included in our analysis of the action phase. To assess if grasp tuning was significant, results were compared to a shuffled condition, where grasp labels were randomly reassigned (see methods).

Tuning was significant during Go-trial peak activity for all brain areas (**Figure 2**C). As expected, tuning was not significant in the ITI condition. During cue phase, results were significant in SMG and PMv, but not significant in S1. During action phase, no significance was found during No-Go trials for all brain areas (**Figure 2**D). These results highlight grasp-dependent neuronal activity during cue presentation in SMG and PMv, and during action phase of Go trials in all brain areas.

How many different grasps was each individual unit able to represent? Results were consistent across all brain areas with the majority of units tuned to one grasp (**Figure 2**E). In SMG and PMv, more units were tuned to multiple grasps, demonstrating mixed grasp encoding within the population. As before, results were significant during cue presentation in SMG and PMv, during Go-trial action phase in all brain areas, but not during ITI, nor during No-Go trial action phase (**Figure 2**E,F).

Similar to previous analysis methodologies (Murata et al., 1997), (Sakata, 1995),(Taira, 1998) (Klaes et al., 2015), we separated tuned units into three categories: those tuned during cue phase (“visual units”), those tuned during Go-trial action phase (“motor-imagery units”) and those tuned during both (“visuo-motor units”). As only SMG and PMv showed significant tuning during the cue phase, this analysis excluded S1 activity. All three neuron types were found. SMG has a higher percentage of units tuned during cue presentation than PMv (75% vs. 58% of all tuned units) which explains a higher overlap of tuned units (47% vs 39%), indicating that cue presentation and motor imagery are more similarly processed in SMG than in PMv (**Figure 2**G).

### SMG, PMv and S1 show significant classification accuracy during grasp motor imagery

To assess each brain region’s potential use for BMI applications, we evaluated decodability of individual imagined grasps using a Naïve bays classification model. A different model was built for each session day. Principal component analysis (PCA) was applied to the dataset (see methods), and leave-one-out classification was performed to compute classification accuracy. To test for significance, results were compared to a null distribution obtained by shuffling the labels before classification (see methods).

Significant motor imagery decoding was observed in all brain areas (**Figure 3**A). Black dots indicate individual session results; red dots indicate averaged shuffled results. Mean +/- 95% confidence interval (c.i.) was computed over individual session. Significant classification accuracies were obtained for cue, delay and Go-action phase in SMG (p < 0.001), cue (p < 0.01), delay (p < 0.05), and Go-action phase (p < 0.001) in PMv, and Go-action phase (p < 0.5) in S1. For No-Go trials, significant classification accuracies were obtained in cue and delay phase for SMG (p < 0.001, p < 0.01), and cue phase in PMv (p < 0.05), but not action phase (**Figure 3**C). Importantly, these results mirror the findings in **Figure 2**C,D, indicating that significant grasp tuning can predict significant classification accuracies. A confusion matrix averaged over all sessions of Go-trials in SMG and PMv during action phase suggests that all grasps can be decoded (**Figure 3**B,D).

**Figure 3.**
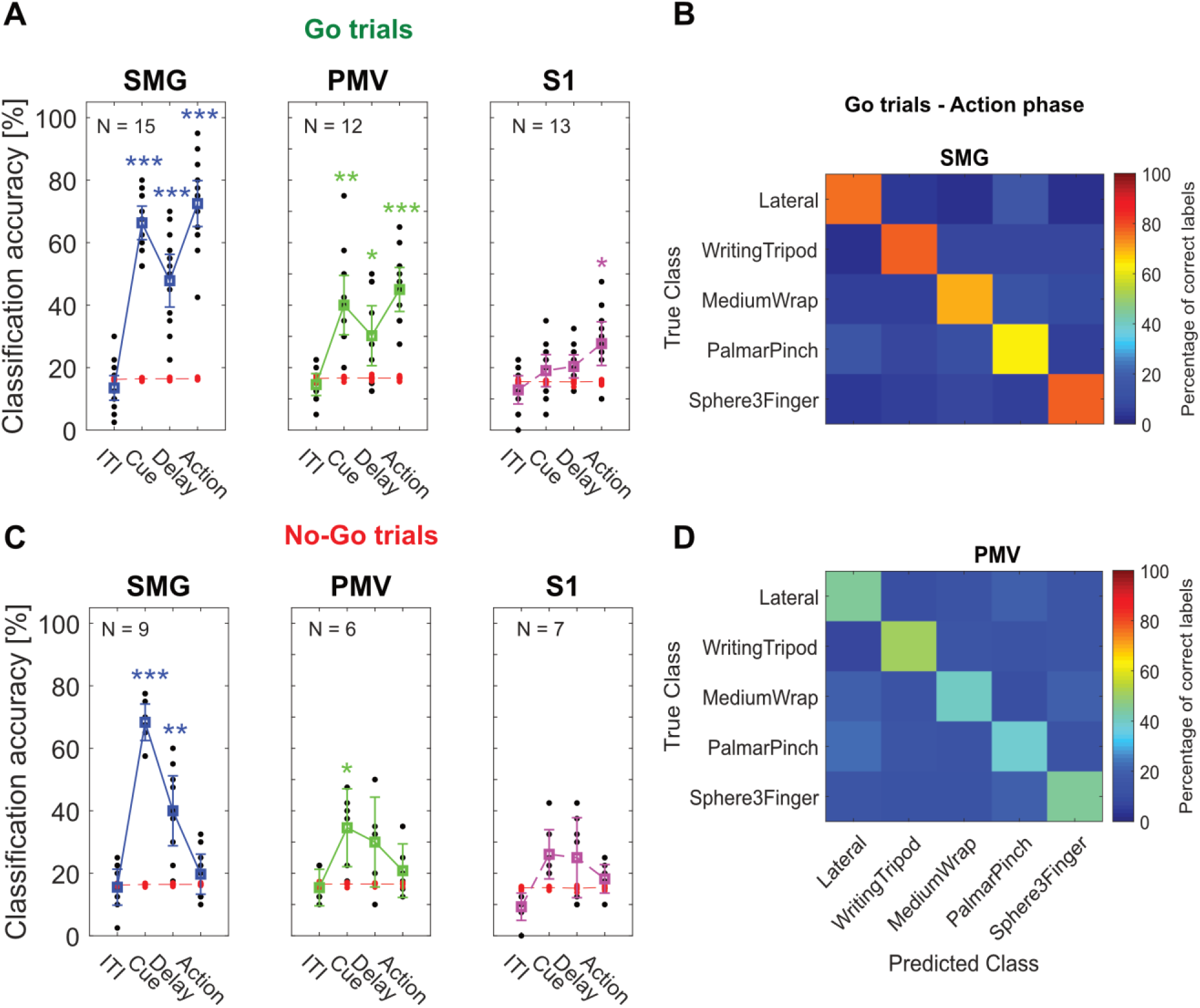
Grasps can be significantly decoded in all brain areas during motor imagery. **A)** Go trials classification accuracy: Classification was performed for each session day individually using leave-one out cross-validation (black dots). 95% c.i. for the session mean was computed. Significance was evaluated by comparing actual data results to a shuffle distribution (averaged shuffle results = red dots, * = p < 0.05, ** = p < 0.01, *** = p < 0.001) **B)** Error matrix during Go-trial action phase for SMG, averaged over all session days. **C)** Same as A) for No-Go trials. **D)** Same as B) for PMV.

### SMG and PMv show high generalizability of grasp encoding in the neural population

We addressed generalizability of grasp encoding in the neural population via two analyses, cross-phase classification and stability across different population sizes.

Cross-phase classification looked at similarities of neural processes across cue and action phases. We trained a classification model on a subset of the data of one phase (e.g. cue phase), and tested it on two different subsets taken from the cue and action phase. If a model trained on the cue phase does not generalize to the action phase, this suggests distinct neural processes are being observed. However, if the model generalizes well, common cognitive processes may be occurring in both phases.

How much could classification accuracy improve if we had a bigger pool of neurons to record from simultaneously? A neuron dropping analysis tracks how classification accuracy evolves as units are removed or added to the pool of predictors. To avoid overfitting, the first 20 principal components were used as features for classification. The analysis was performed separately for each of the implanted brain regions, over 100 repetitions of eight-fold cross-validation. As explained previously, one dataset was compiled from training on cue phase and evaluating on both cue and action phase, and another dataset from training on the action phase, and evaluating over both cue and action phases (see methods).

Results of both analyses are represented in **Figure 4**. SMG and PMv show strong shared activity between cue and action phase. When training on cue phase, and testing on cue and action phase, we observe that the model generalizes very well in SMG, with overlapping 95% bootstrapped confidence intervals that diverge only at high unit count. In PMv, the generalization is a bit lower, but shows similar trends, while decoding remains at chance level for S1 (**Figure 4**A,B,C Train: Cue Phase). However, when training on the action phase, and evaluating on the cue phase, lower generalization of the model is observed in SMG and PMv (**Figure 4** A,B,C Train: Action Phase).

**Figure 4.**
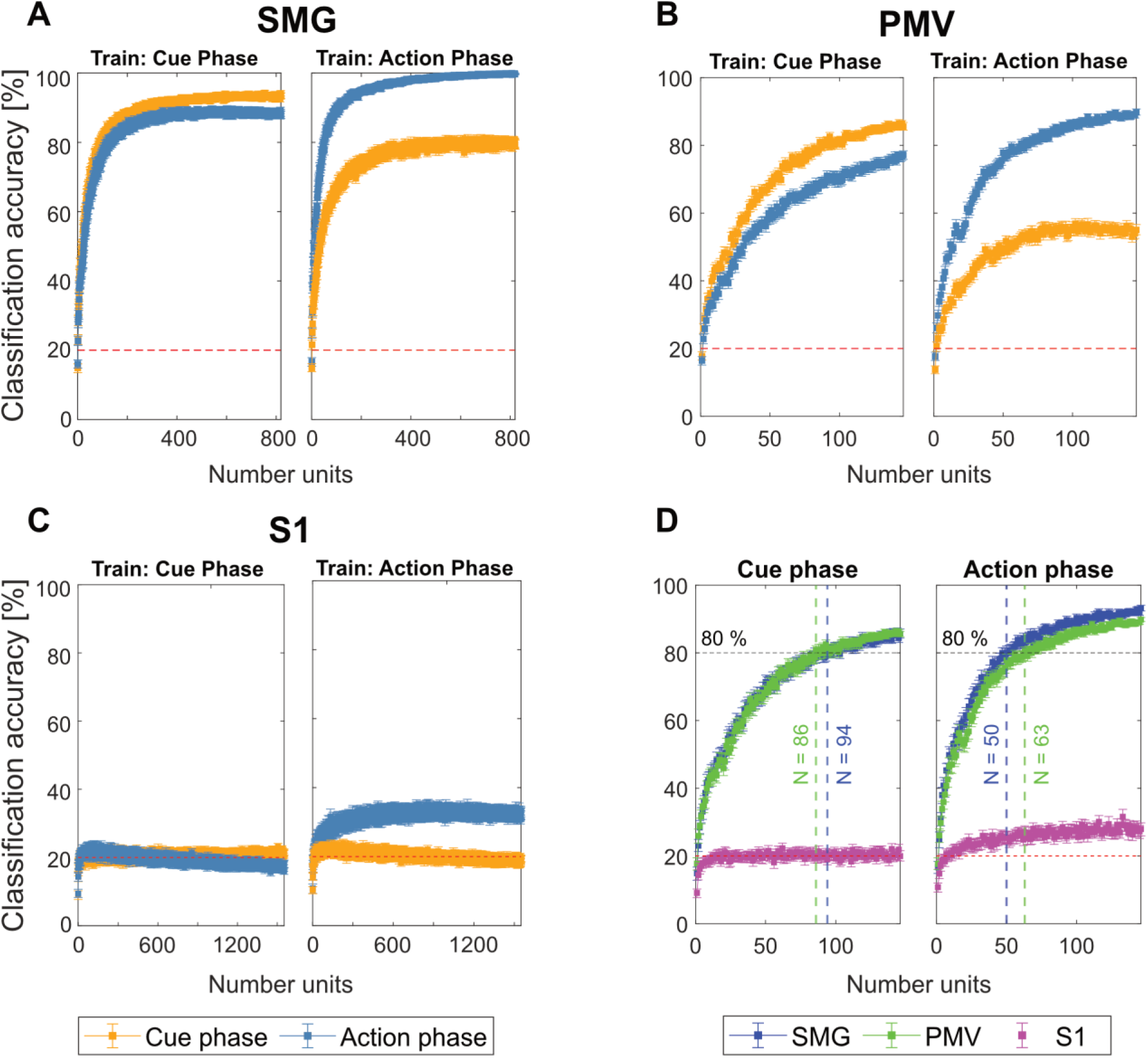
SMG and PMV show high generalizability of grasp encoding in neuronal populations. **A-C)** A neuron dropping curve was performed in SMG, PMv and S1 over 100 repetitions of eight-fold cross validation. The analysis was performed once by training the model on the cue phase and applying it on both cue and action phase (Train: Cue phase), and once by training it on the action phase and applying it on both cue and action phase (Train: Action phase). The mean classification accuracy with bootstrapped 95% c.i. was plotted. **D)** To compare decoding performance between different brain areas, the first 140 units of each brain areas were plotted together. The number of units needed to obtain 80% classification accuracy during cue phase and action phase was calculated. SMG and PMv results were similar, with less units needed for classification during action phase compared to cue phase.

During the action phase, SMG peaks at 99% decoding accuracy when all recorded units are included in the analysis (**Figure 4**A). In S1, decoding accuracy during action phase peaks around 32%, even when the pool of available neurons increases (**Figure 4**C). As PMv did not reach its peak decoding accuracy due to fewer number of units recorded (**Figure 4**B), performance of SMG and PMv at same population levels was compared directly. **Figure 4**D depicts the number of features needed to obtain 80% classification accuracy during cue (left) and action (right) phase. During cue phase, 94 units in SMG and 86 units in PMv were needed. During action phase, 80% classification accuracy was obtained with 50 units in SMG, and 63 units in PMv. These results demonstrate that SMG’s and PMv’s potential for the decoding of grasps is comparable, and that if a higher neuronal population were available to record from simultaneously, excellent grasp classification results can be expected in both brain areas.

### SMG significantly decodes spoken grasps and colors

To explore SMG, PMv and S1’s role in a different cognitive processing task, the participant was instructed to perform verbal speech instead of motor imagery during action phase. By comparing each region’s evoked activity between these two cognitive processes, we aimed to find evidence for language processing activity at the single unit level. During each session day, a “Motor Imagery” (or Go task, see methods), a “Spoken Grasps” and a “Spoken Colors” version of the task was run. During the “Motor Imagery”, grasp motor imagery was performed during action phase. In the “Spoken Grasps” version of the task, the participant was instructed to say aloud the name of the visually cued grasp, instead of performing motor imagery during action phase. In the “Spoken Colors” version of the task, the participant was cued with visual depictions of colors, and said aloud the name of the color during the action phase (**Figure 5**A,B).

**Figure 5.**
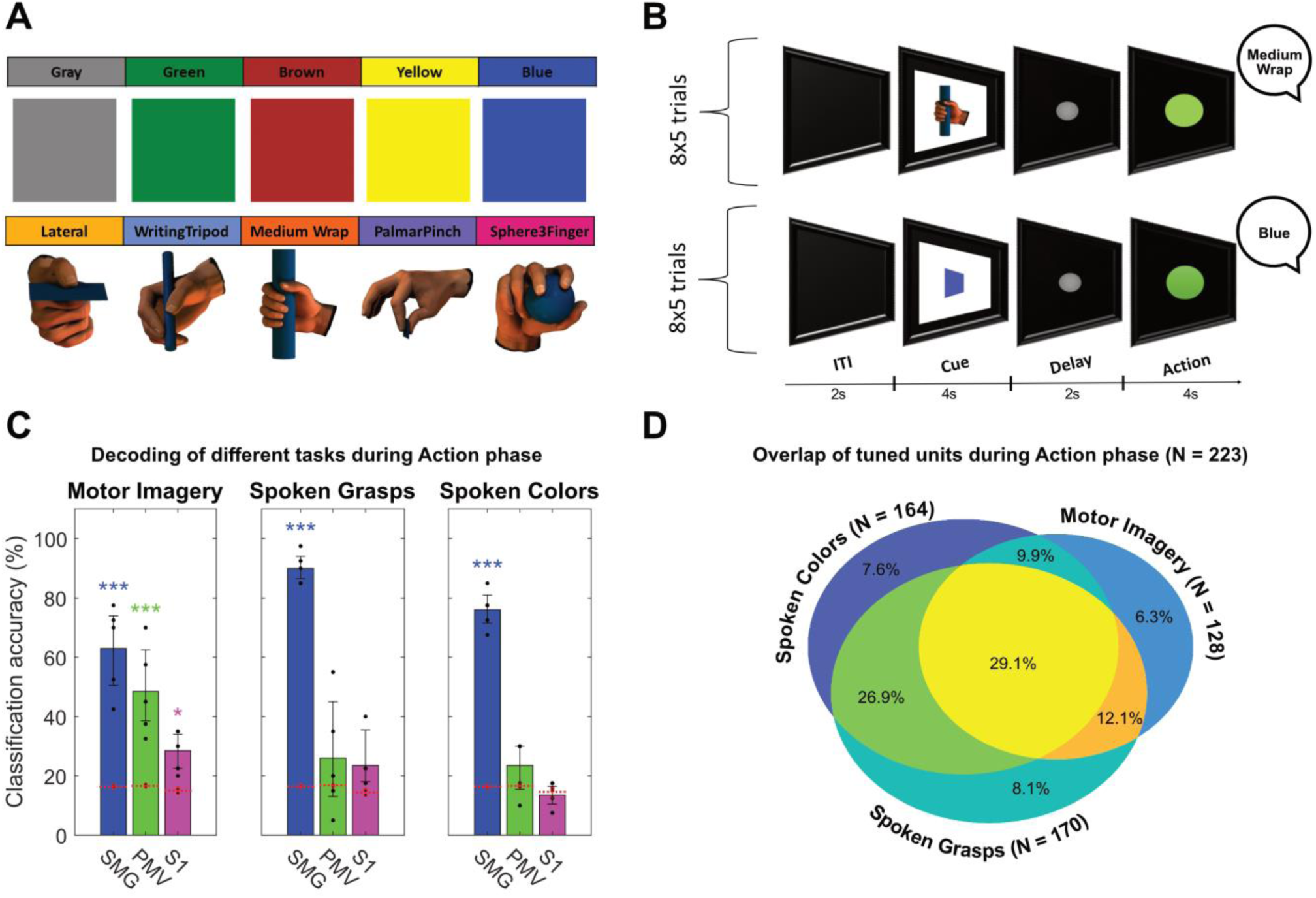
SMG encodes speech. A control task was performed, where speech instead of motor imagery was studied during the action phase. **A)** Grasps images and color images were used to cue speech in a tetraplegic human. **B)** The task was composed of an inter-trial interval (ITI), a cue phase displaying the image of one of the grasp or colors, a delay phase and an action phase. During the action phase, the participant was instructed to say out loud the name of the cued grasp or color. **C)** Classification was performed for each session day individually using leave-one out cross-validation (black dots). 95% c.i. for the session mean was computed. Results during the action phase are shown. SMG, PMv, and S1 showed significant classification results when motor imagery was performed. Only SMG showed significant classification results during spoken grasps and spoken colors. **D)** Overlap of tuned channel during the action phase between “Motor Imagery”, “Spoken Grasps” and “Spoken Colors” task. While most units were tuned during all tasks, (29.1%), there was a higher overlap of units tuned both during color and grasp speech (26.9%), than during grasp speech and grasp motor imagery (12.1%), and color speech and grasp motor imagery (9.9%).

Classification results during the action phase corroborate SMG’s involvement during language processing (**Figure 5**C) (Oberhuber et al., 2016)(Deschamps et al., 2014) (Stoeckel et al., 2009). During motor imagery, SMG, PMv and S1 show significant classification accuracy. However, during speech production, only SMG shows significant results, both for spoken grasp names, and spoken colors.

We inspected if units in SMG were tuned (using linear regression) during motor imagery, speech of grasps, and/or speech of colors, and represented the results in a Venn diagram (**Figure 5**D). This analysis allows us to probe if the output modality (speech vs. motor imagery), or the semantic content (grasps vs. colors) was more similarly represented in the neuronal data. While most units were tuned during the three tasks (29.1%), a higher overlap of tuned units was shown between the speech conditions (26.9%), than between speech and motor imagery condition (12.1%). These results suggest that the output modality is more similarly encoded than the semantic content.

## Discussion

In this work, we demonstrated that motor imagery of five unique grasps was well represented by the firing rates of neuronal populations, and could be decoded significantly above chance level in the supramarginal gyrus (SMG), the ventral premotor cortex (PMv), and the primary sensory cortex (S1). SMG and PMv encoded grasp information both during cue presentation and during motor imagery with similar neuronal activity patterns. Equal numbers of units in the neuronal populations of SMG and PMv showed comparably excellent grasp encoding capabilities, demonstrating high potential for grasp BMI applications in both areas. Spoken names of five grasps and five colors were decodable in SMG, indicating a potential site for speech BMI synthesizer applications.

To demonstrate the participant had volitional control of motor imagery during the action phase, and observed activity was not due to some external factor, interleaved No-Go trials served as a control. During No-Go trials in the action phase, unit tuning was not significantly different from a shuffled distribution (**Figure 2**D,F), and classification was not significantly different from chance (**Figure 3**A). While a non-significant peak in tuning was observed in **Figure 2**B (No-Go trials action), this activity likely represented grasp-independent modulation of the firing rate. As the participant was only instructed to perform or abort motor imagery at the beginning of the action phase, formation of a motor plan might have already occurred, requiring a certain processing times to cancel. Similar effects have been observed in PPC when choosing one effector (saccade or reaches), while the other get canceled (Cui and Andersen, 2007). However, as different grasps are not decodable during No-Go action phase (**Figure 3**A, No-Go trials action phase), it might be evidence for the participant actively canceling motor plans, an important feature for a grasp BMI applications.

### S1 encodes imagined grasps significantly, but does not improve with population size

Decoding grasp intentions in the same brain area where sensations can be elicited would allow for the design of a bidirectional BMI device that requires fewer implantation sites. Microstimulation of this participants’ S1 array resulted in sensations over arm and hand area (Armenta Salas et al., 2018). In our work, S1 grasp motor imagery classification was significant (**Figure 3**A). However, performance did not improve with increased population sizes as much as SMG and PMv (**Figure 4**D). Several reasons might explain this behavior. Firstly, no actual movement was performed, likely decreasing the occurrence of proprioceptive signals. Secondly, the task design might have only weakly engaged the neural populations we recorded from, as the electrode implant mostly covered the contralateral arm area (Armenta Salas et al., 2018). A different task, that involved the arm by reaching to grasp an object, could have elicited stronger neuronal activity (Jafari et al., 2020). Thirdly, units in S1 showed mostly grasp independent increase in activity compared to baseline (**Figure 2**C,E), opening up the possibility that the grasps were not different enough to evoke stronger decoding abilities in S1.

### SMG and PMv show significant activity during visual cue presentation of grasps

SMG demonstrated a faster rise of activity at cue presentation than PMv (**Figure 2**A,B). Similar results between PPC and ventral premotor cortex have been shown in NHP (Schaffelhofer and Scherberger, 2016). This observation supports the hypothesis that SMG relays planning information to PMv through anatomical connections (Ramayya et al., 2010),(Koch et al., 2010). We found units encoding each of the five grasps (**Figure 2**A). Furthermore, we identified units that were only active during cue phase (visual), units only active during action phase (motor), and units active during both (visuo-motor), as seen in NHP and human studies (Akira Murata et al., 2000), (Carpaneto et al., 2011),(Klaes et al., 2015),(Schaffelhofer et al., 2015) (**Figure 2**G).

### Evidence for mixed visual and motor activity during action phase

While human participants can self-report strategies employed while performing internal cognitive tasks, cue processing and motor imagery do not have independently observable behavioral outputs. Therefore, multiple explanations for synchronized neural activity observed during these tasks are plausible. During cue presentation, an increase in neural activity could represent visual feature extraction of the presented cue (visual processes). Alternatively, activity could be independent of visual input and represent planning activity of the cued grasp (motor processes). Additionally, activity could be memory or semantic, as the participant remembers the instructed grasp (cognitive processes). Finally, a combination of all these processes might be at play. While proving a definitive answer to this question is beyond the scope of this paper, performing cross-phase classification between the cue and action can help identify similar or distinct cognitive processes within the observed data.

Cross-phase classification found similar neuronal activity in cue and action phase in both SMG and PMv (**Figure 4**A,B). This agrees with our finding of overlapping neuronal populations tuned during both cue and action phase (**Figure 2**G). One explanation for these similarities could be the participant performing “visual imagery” rather than motor imagery during action phase, by recalling a mental image of the grasp (**Figure 4**A,B Train: Cue Phase). Cue phase activity can partly be explained during action phase (classification performance 80% SMG, 55% PMv) (**Figure 4**A,B Train: Action Phase), but neuronal activity unique to action phase exists (classification performance 99% SMG, 89% PMv). This generalization from the cue to action phase is not bidirectional (from action to cue phase), therefore we argue this additional information during the action phase is likely motor.

Good generalization of the model to both phases when training on the cue phase could indicate that cue phase activity has motor components as well as visual. While AIP is assumed to extract visual features of objects during cue and delay phase, PMv has been speculated to represent planning activity of the grasp. Neurons in NHP ventral premotor area F5 have demonstrated higher neuronal modulation during cue and planning phase when different objects were grasped with different grasps, rather than different objects grasped with the same grasp. This evidence demonstrates that the planned hand shape, and not only visual object features, can modulate neuronal firing during cue and delay phase in the grasp circuit (Schaffelhofer and Scherberger, 2016). Similar effects might be at play in our observed data.

Cue phase activity could represent semantic or memory processing, i.e. the abstract concept of each cued grasp. During tool use, SMG is hypothesized to integrate the appropriate grasp type with the knowledge of how to use the tool (Osiurak and Badets, 2016; Vingerhoets, 2014), which requires access to semantic information. As our current task design does not allow the differentiations of these cognitive processes, further experimentation is necessary. For instance, cueing grasps with non-visual sensory cues and observing if cue phase activity is still present, might allow the dissociation between visual, motor and semantic processes, and help clarify the roles of SMG and PMv in the human grasp circuit.

When analyzing SMG and PMv for their potential for grasp BMI applications, both performed similarly. While SMG displays stronger encoding of grasps than PMv on a session-to-session basis (**Figure 3**A), these results are likely due to the small number of units we were able to record from the PMv array on individual days. The neuron dropping analysis illustrates when identical neuronal population are present, SMG and PMv have similar decoding abilities (**Figure 4**D).

### SMG encodes speech

During speech, SMG and PMv showed vastly different results. Spoken words (both grasp names and colors) were decodable equally or better than only motor imagery of grasps in SMG. In contrast, PMv and S1 showed neither significant classification of spoken grasp names nor spoken colors (**Figure 5**C). These results could indicate that SMG processes semantics, regardless of the performed task. However, in a different analysis (**Figure 5**D), neuronal population tuning was more similar for output modality (speaking of colors and speaking of grasps) than for semantics (motor imagery of grasps and speaking of grasps), hinting that other cognitive processes are at play. As spoken color names were also decodable, SMG’s role is not confined to only action verbs. These results are evidence for a larger role of SMG in language processing, and indicate SMG as a candidate implant site for a speech BMI (Andersen et al., 2014).

## Conclusion

In this paper, we demonstrated grasps were well represented by firing rates of neuronal populations in human SMG and PMv during cue presentation. During motor imagery, individual grasps could be significantly decoded in all brain areas. SMG and PMv achieved similar highly-significant decoding performances, demonstrating their viability for a grasp BMI. During speech, SMG achieved significant classification performance, in contrast to PMv and S1, which were not able to significantly decode individual spoken words.

These findings suggest that grasp signals can be robustly decoded at a single unit level from the cortical grasping circuit in human. Our data suggests PMv has a specialized role in grasping, while SMG’s role is broader and extends to language processing. Together, these results indicate that brain signals from high-level areas of the human cortex can be exploited for a variety of different BMI applications, and further demonstrate the potential for BMIs to provide increased independence for people living with tetraplegia.

## Acknowledgements

We wish to thank L. Bashford, H. Jo and I. Rosenthal for helpful discussions and data collection. We wish to thank our study participant FG for the continuous contributions that made this work possible.

